# Longitudinal prediction of outcome in idiopathic pulmonary fibrosis using automated CT analysis

**DOI:** 10.1101/493544

**Authors:** Joseph Jacob, Brian J. Bartholmai, Coline H.M. van Moorsel, Srinivasan Rajagopalan, Anand Devaraj, Hendrik W. van Es, Teng Moua, Frouke T. van Beek, Ryan Clay, Marcel Veltkamp, Maria Kokosi, Angelo de Lauretis, Eoin P. Judge, Teresa Burd, Tobias Peikert, Ronald Karwoski, Fabien Maldonado, Elisabetta Renzoni, Toby M. Maher, Andre Altmann, Athol U. Wells

## Abstract

**AIMS:** To evaluate computer-derived (CALIPER) CT variables against FVC change as potential drug trials endpoints in IPF.

**METHODS:** 71 Royal Brompton Hospital (discovery cohort) and 23 Mayo Clinic Rochester and 24 St Antonius Hospital Nieuwegein IPF patients (validation cohort) were analysed. Patients had two CTs performed 5-30 months apart, concurrent FVC measurements and were not exposed to antifibrotics (to avoid confounding of mortality relationships from antifibrotic use). Cox regression analyses (adjusted for patient age and gender) evaluated outcome for annualized FVC and CALIPER vessel-related structures (VRS) change and examined the added prognostic value of thresholded VRS changes beyond standard FVC change thresholds.

**RESULTS:** Change in VRS was a stronger outcome predictor than FVC decline when examined as continuous variables, in discovery and validation cohorts. When FVC decline (≥10%) and VRS thresholds were examined together, the majority of VRS change thresholds independently predicted outcome, with no decrease in model fit. When analysed as co-endpoints, a VRS threshold of ≥0·40 identified 30% more patients reaching an endpoint than a ≥10% FVC decline threshold alone.

**CONCLUSIONS:** Change in VRS is a strong predictor of outcome in IPF and can increase power in future drug trials when used as a co-endpoint alongside FVC change.

**Ethics committee approval:** Approval for this study of clinically indicated CT and pulmonary function data was obtained from Liverpool Research Ethics Committee (Reference: 14/NW/0028) and the Institutional Ethics Committee of the Royal Brompton Hospital, Mayo Clinic Rochester and St. Antonius Hospital, Nieuwegein. Informed patient consent was not required.

**Take home message:** Change in the vessel-related structures, a computer-derived CT variable, is a strong predictor of outcome in idiopathic pulmonary fibrosis and can increase power in future drug trials when used as a co-endpoint alongside forced vital capacity change.

## INTRODUCTION

The advent of antifibrotic agents ^1,2^ has spelled the end of the placebo-controlled trial in idiopathic pulmonary fibrosis (IPF). Validation of new therapies against nintedanib or pirfenidone will need to identify smaller increments of improvement when compared to placebo-controlled trials. The current gold standard endpoint in IPF drug trials is change in forced vital capacity (FVC). However, the measurement variation associated with FVC testing risks masking genuine beneficial effects from a new drug in a trial setting^3^. More sensitive and accurate endpoints are required for IPF drug trials.

Computer advances have resulted in the development of new tools capable of quantifying disease extents on CT with excellent reproducibility and precision. Modern computer algorithms can quantify CT patterns with clear visual analogues as well as patterns that may not be easily visually recognized. An example is the quantitation of pulmonary vessel-related structures (CAL VRS) by CALIPER which has been shown to powerfully predict outcome at baseline in IPF^4^.

The primary aim of the current study was to evaluate whether quantitative changes in computer features across serial CT examinations can predict mortality in independent discovery and validation populations of IPF patients. We secondarily evaluated thresholds of change in computer-derived variables as potential adjunctive endpoints in patients with equivocal FVC change.

## METHODS

### Study Design

IPF was diagnosed by multidisciplinary teams in patients receiving two non-contrast volumetric CT scans between 5 and 30 months apart as part of their clinical care. Previous baseline analyses of IPF patients^5^ made it apparent that variable initiation time of antifibrotics and varied dosages, durations and types of antifibrotic medication in study participants had a profound confounding effect on mortality relationships. Specifically, patients not uncommonly began antifibrotics between the first and second CTs and in most cases after the second CT. Consequently, cardinal analyses in the current manuscript were restricted to patients not receiving any anti-fibrotic therapy (n=118).

The discovery cohort comprised 71 consecutive IPF patients presenting to the Royal Brompton Hospital between January 2007 and December 2014. The validation cohort comprised IPF patients presenting to either the St Antonius Hospital, Nieuwegein (n=24) between January 2005 and June 2014 or the Mayo Clinic Rochester (n=23) between January 2009 and June 2015.

Pulmonary function tests analysed at baseline included FVC and diffusion capacity of carbon monoxide (DLco). FVC examined longitudinally was restricted to FVC collected within three months of the second CT to allow for a fair comparison between CT-derived variables and FVC. Pulmonary function test and CT protocols are outlined in the Supplementary Appendix.

### CT pattern evaluation

27 CT features were scored by CALIPER. Nine features were measured on a whole lung level and included total lung volume, normal parenchyma, emphysema, honeycombing, reticular pattern, ground glass opacity and CAL VRS (details in Supplementary appendix). Fibrosis extent represented the sum of reticular pattern and honeycombing. Interstitial lung disease (ILD) extent additionally summed ground-glass opacification.

In addition to the 9 whole lung features, 18 CAL VRS subdivisions were evaluated. Separation was according to the lung zonal location of the structures: upper (UZ), middle (MZ) and lower zones (LZ), and the cross-sectional area of the structures in each zone: <5mm^2^, 5-10mm^2^, 10-15mm^2^, 15-20mm^2^, >20mm^2^. Volumes for all CALIPER features were converted into a percentage using CALIPER-derived total lung volume measurements^4,6^. Absolute change in the derived 27 CT variables was calculated by dividing the absolute difference (CT2-CT1) by the time interval between the two measurements (in years). CALIPER CT analysis methods are detailed in the Supplementary appendix.

### Statistical Analyses

#### Demographics

Data are given as means with standard deviations, or patients numbers with percentages where appropriate. Mean group differences were evaluated using a Chi-squared test for categorical variables, a two-sample T test for parametric continuous variables, and the Mann-Whitney U test for medians. The McNemar test compared patient numbers reaching different study endpoints. Pearson’s correlation examined linkages between FVC change and VRS/UZ VRS change.

#### Derived variables

Annualised FVC change was measured using two different methods: a linear mixed effects model on all eligible timepoints and a naïve estimate from only two timepoints (first and last). For the naïve estimate we computed annual relative change by dividing the absolute annual change by the baseline FVC value (relative). As a more comprehensive estimate of FVC change incorporating all eligible timepoints we used linear mixed effect models to derive the best linear unbiased predictor (BLUP). The lmer function from the R package lme4^7^ was used for the analysis. From the BLUP slopes we derived the annualized change relative to the first FVC measurement. In addition, to the quantitative annualized decline we derived dichotomised declines indicating whether there was a ≥5% or ≥10% reduction in FVC compared to baseline value based on the BLUP estimate. Finally, we also extracted the FVC measurement closest to the second CT scan to indicate the added value of longitudinal measurements over single timepoints.

We analysed clinical progression using Cox proportional hazards models, and evaluated model fit using C-indices. Time was measured from the timepoint of the second CT and an event was either death (n=90) or transplantation (n=8). Each predictor variable was tested alone while correcting for confounders (age at the second CT scan and sex). For completeness, a secondary analysis on all subjects regardless of their history of antifibrotics usage is included in the supplement (n=104 discovery cohort; n=96 validation cohort).

#### Examining vessel score thresholds

As clinical progression is often measured using dichotomised response rather than quantitative response, we examined survival using thresholds of CAL VRS and UZ VRS change and measured these against relative FVC decline thresholds of ≥5% and ≥10% derived from the BLUP estimates. More precisely, we used the absolute change in CAL VRS and UZ VRS and created a series of binary variables indicating an increase of more than 0.0, 0.1, 0.2, …1.0 for both VRS variables. We tested for the significance of these indicators in the presence of either ≥5% or ≥10% FVC decline indicator variables and covariates (age at second CT and sex) in a Cox proportional hazards model (n=118). The corresponding p-value indicates the improvement in model fit with the additional information. We also carried out the reverse analysis, i.e., testing the p-value for either ≥5% or ≥10% decline when added to a model containing an indicator variable for VRS decline.

We also compared the predictive performance of FVC-based indicator variables to VRS indicator variables, either used alone or when combined with an FVC-based indicator variable as a ‘joint endpoint’. The joint endpoint reflected whether the FVC decline or the VRS increase was achieved with estimates based on 500 bootstrap replicates (n=118). We estimated the number of additional patients that would reach either a ≥5% or ≥10% predicted FVC threshold or a preselected CAL VRS/UZ VRS threshold in a drug trial setting. Further, we computed the Kaplan-Meier estimator for different subgroups of patients (n=118) depending on whether they reached the FVC or VRS threshold or both (using the SPSS Kaplan Meier function^11^). Finally, we examined mortality prediction in patients with a BLUP estimated relative decline in FVC of >5% but <10% not receiving antifibrotics (n=41). Cox proportional hazards models examined the thresholded CAL VRS and UZ VRS variables and computed the C-index based on 500 bootstrap replicates.

## RESULTS

### Demographic Data

All CTs in the discovery cohort (n=71) were acquired using a B70 Siemens kernel. CTs in the validation cohort (n=47) were acquired using six different kernels and in 14/118 (12%) patients, CT kernels varied across the two study CTs. Baseline differences between the two cohorts are shown in Supplementary Table 1. No patients were lost to follow up. 33/104 (32%) patients in the discovery cohort and 49/96 (51%) patients in the validation cohort (p=0.006) receiving antifibrotics were excluded from the cardinal analysis. The mean CT interval was similar for discovery (1.1 years) and validation (1.2 years) cohorts. Significant but weak correlations (r=-0.42, p=1.8×10^-6^) were identified between FVC change (measured using BLUP estimates) and absolute VRS change (Supplementary Table 2).

**Table 1.** C-indices demonstrating model fit for mortality in IPF patients not exposed to antifibrotic medication (top) and IPF patients regardless of exposure to antifibrotic medication. FVC=forced vital capacity, VRS=vessel-related structures.

| **Antifibrotic Use** | **Variable**<br>**(Units are percentage**<br>**unless specified)** | **Discovery**<br>**Cohort**<br>**(C-index)** | **Validation**<br>**Cohort**<br>**(C-index)** | **Combined**<br>**Population**<br>**(C-index)** |
| --- | --- | --- | --- | --- |
| Patients without<br>antifibrotic<br>exposure | CALIPER VRS change | 0.64 | 0.62 | 0.64 |
|  | Upper zone VRS change | 0.65 | 0.63 | 0.64 |
|  | Relative FVC change | 0.65 | 0.63 | 0.65 |
| Patients with and<br>without antifibrotic<br>exposure | CALIPER VRS change | 0.65 | 0.72 | 0.69 |
|  | Upper zone VRS change | 0.65 | 0.71 | 0.68 |
|  | Relative FVC change | 0.65 | 0.63 | 0.66 |

**Table 2.**
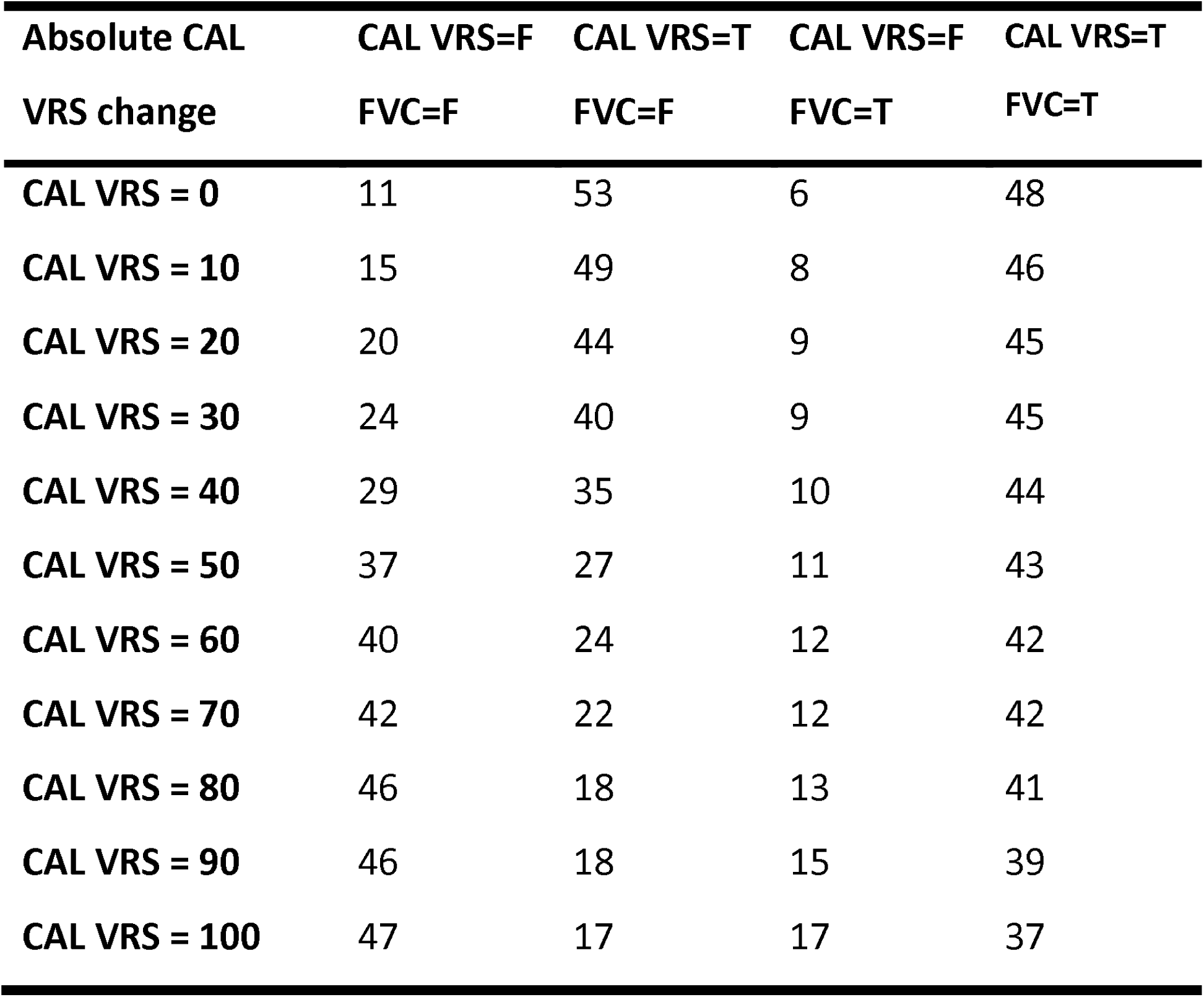
Numbers of patients reaching threshold endpoints of CALIPER vessel-related structure increase (CAL VRS) or ≥10% relative forced vital capacity (FVC) decline from the study population of 118 patients not exposed to antifibrotic medication. T=true (did reach threshold), F=false (did not reach threshold).

| **Absolute CAL**<br>**VRS change** | **CAL VRS=F**<br>**FVC=F** | **CAL VRS=T**<br>**FVC=F** | **CAL VRS=F**<br>**FVC=T** | **CAL VRS=T**<br>**FVC=T** |
| --- | --- | --- | --- | --- |
| **CAL VRS = 0** | 11 | 53 | 6 | 48 |
| **CAL VRS = 10** | 15 | 49 | 8 | 46 |
| **CAL VRS = 20** | 20 | 44 | 9 | 45 |
| **CAL VRS = 30** | 24 | 40 | 9 | 45 |
| **CAL VRS = 40** | 29 | 35 | 10 | 44 |
| **CAL VRS = 50** | 37 | 27 | 11 | 43 |
| **CAL VRS = 60** | 40 | 24 | 12 | 42 |
| **CAL VRS = 70** | 42 | 22 | 12 | 42 |
| **CAL VRS = 80** | 46 | 18 | 13 | 41 |
| **CAL VRS = 90** | 46 | 18 | 15 | 39 |
| **CAL VRS = 100** | 47 | 17 | 17 | 37 |

### Discovery and validation cohort mortality analyses

CAL VRS and UZ VRS measures were the strongest predictors of outcome in discovery and validation cohorts (Figure 1) and were at least equivalent to FVC change when evaluated using C-indices (Table 1). Longitudinal measures of either FVC decline or CT scores provided stronger prognostic information than the single FVC measure closest to the second CT scan. The effects of confounding from antifibrotic use on mortality in all IPF patients in the discovery and validation populations regardless of antifibrotic use are shown in Supplementary Figure 1.

**Figure 1.**
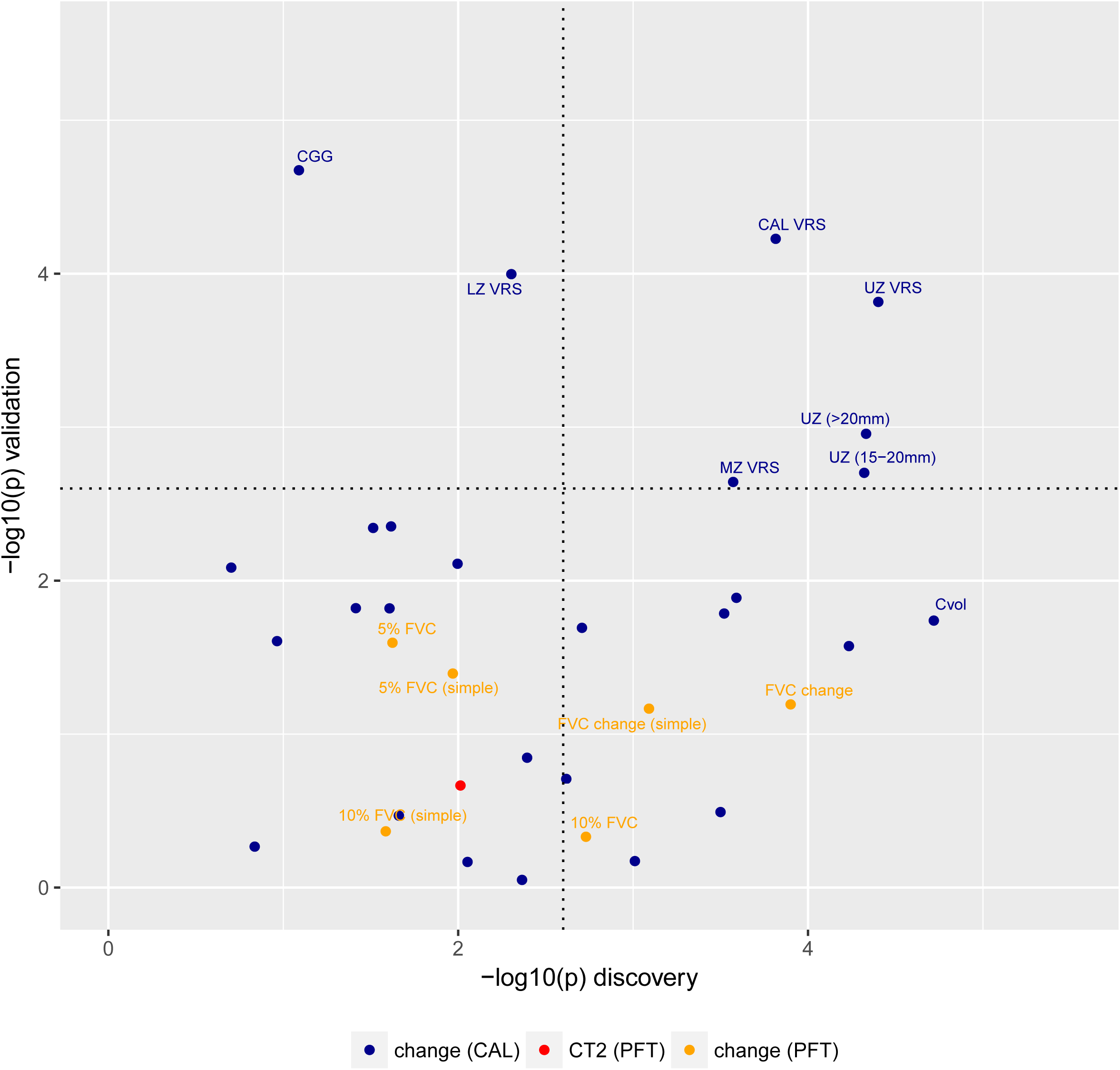
Scatterplots demonstrating -log10 p-values for various computer-derived (CALIPER) variables (blue points) and FVC decline (orange points) in patients not exposed to antifibrotic medication (Figure 1) in the discovery cohort (x-axis, n=71) and validation cohort (y-axis, n=47). Horizontal and vertical dotted lines represent the Li and Ji corrected cutoff for statistical significance. FVC decline was calculated using two methods: naïve estimate from two timepoints aligned with the two CT timepoints (simple) and using best linear unbiased predictions. FVC change was expressed as a continuous variable (FVC change), and at ≥5% decline and ≥10% decline thresholds. The FVC value at the timepoint of the second CT scan (red dot) was used to benchmark expressions of FVC decline. The pulmonary vessel-related structure score (CAL VRS) was subdivided according to zonal location (UZ VRS=upper zone, MZ VRS=middle zone, LZ VRS=lower zone) and structure cross-sectional area in each zone (<5mm^2^, 5-10 mm^2^, 10-15 mm^2^, 15-20 mm^2^, >20 mm^2^).

### VRS Threshold Mortality Analyses

Multivariate Cox mortality models were used to evaluate thresholds of change in CAL VRS/UZ VRS against relative ≥5% or ≥10% FVC decline thresholds in the combined population of patients not receiving antifibrotics (n=118). A CAL VRS threshold of ≥0.30 independently predicted mortality when evaluated against a ≥10% FVC decline threshold (Figure 2a), but at CAL VRS thresholds of ≥0.50, a ≥10% FVC decline threshold no longer significantly contributed to predicting mortality (Figure 2b). Similar trends were seen when UZ VRS was examined against a ≥10% FVC decline threshold and when both CALIPER variables were examined against ≥5% FVC decline thresholds (Figure 2a+b). Overall, more severe changes in VRS variables (e.g., ≥0.50) significantly improved the model fit over FVC variables alone (Figure 2a).

**Figure 2.**
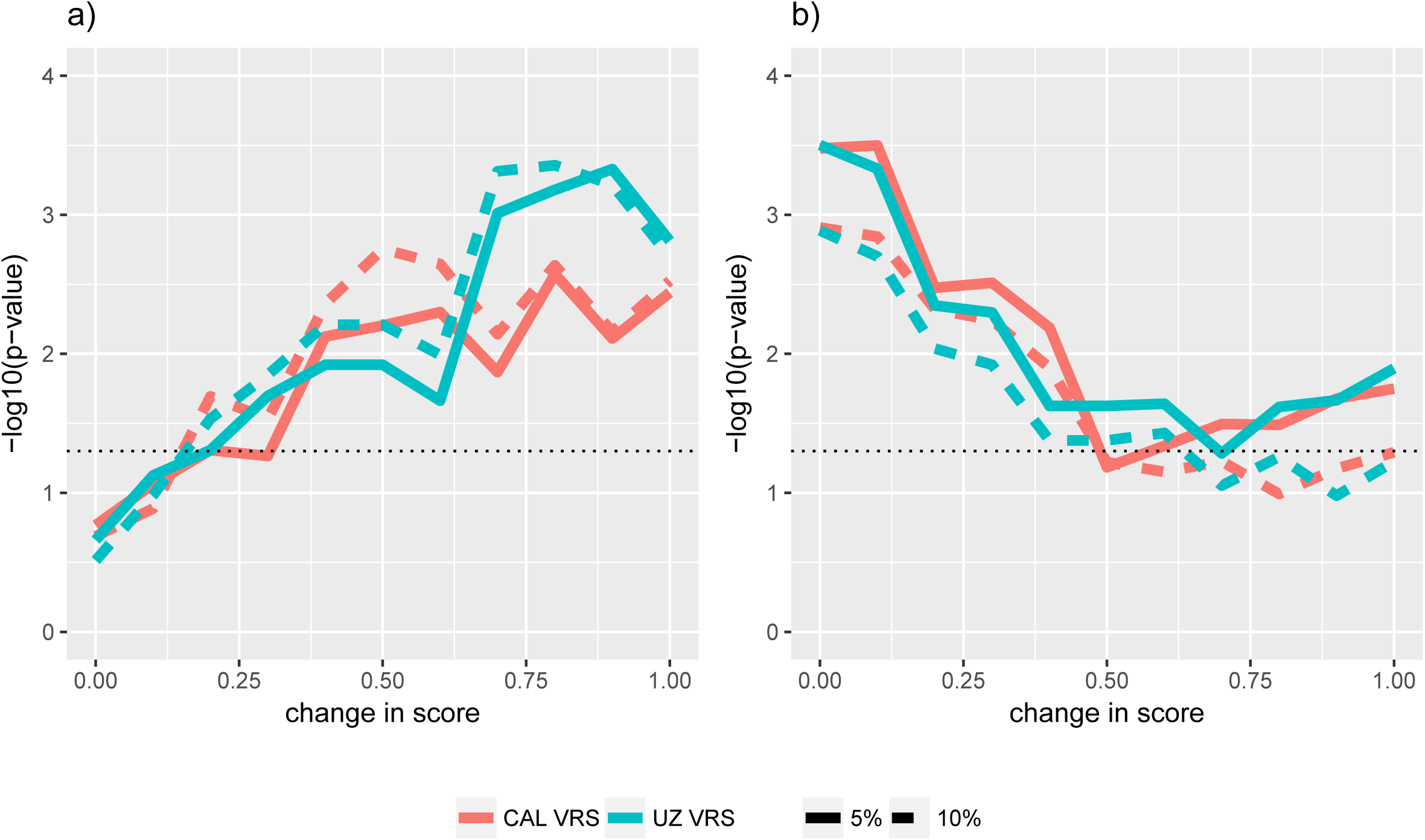
Line graphs demonstrating -log10 p-values (y-axis) for covariates in multivariate Cox mortality models examining change at varying thresholds (x-axis) in vessel related structures (CAL VRS, red) and upper-zone vessel related structures (UZ VRS, green). (a) P-values for models where CAL VRS and UZ VRS thresholded variables have been added to models containing a ≥5% FVC decline threshold (solid line) or a ≥10% FVC decline threshold (dashed line). (b) P-values for the ≥5% (solid) and ≥10% FVC (dashed) decline thresholds when added to Cox models containing the thresholded CAL VRS (red) or UZ VRS (green) variables. A horizontal black dotted line indicates the unadjusted 0.05 cut-off for statistical significance.

At CAL VRS and UZ VRS thresholds of ≥0.40, no difference in model C-index was seen when compared to a ≥10% FVC decline threshold (Figure 3a+b) The C-index was unchanged when using either a solitary CALIPER endpoint (CAL VRS or UZ VRS), or a combined endpoint of a CALIPER variable and an FVC ≥10% decline threshold. Results were maintained when CALIPER variable thresholds were compared to a ≥5% FVC decline threshold (Figure 3c+d).

**Figure 3.**
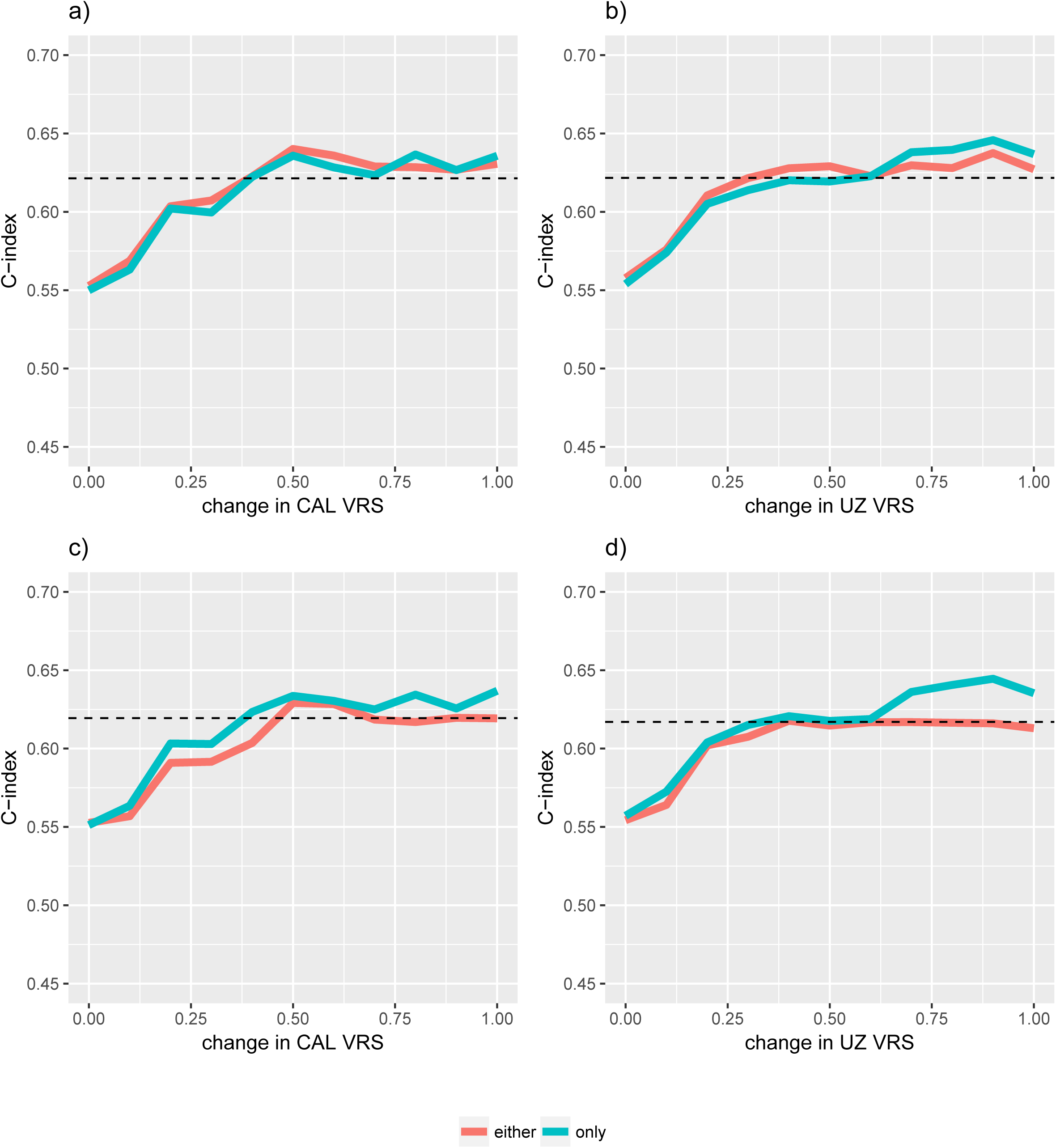
Graphs demonstrating C-indices for models examining varying thresholds of change in vessel related structures (CAL VRS, panels a+c) and upper-zone vessel related structures (UZ VRS, panels b+d). The CAL VRS and UZ VRS thresholds have been separately examined against either a 10% FVC decline threshold (panels a-b) or a 5% FVC decline threshold (panels c-d). The horizontal dotted black line indicates the C-index for models examining the relevant FVC threshold alone. Green lines demonstrate the C-indices for models when a CALIPER (CAL VRS or UZ VRS) threshold alone was examined. Red lines demonstrate the C-indices for models where a binary variable indicated a ‘joint endpoint’, i.e., either the CALIPER or FVC threshold had been reached.

### Sensitivity of FVC and VRS Thresholds

79/118 (67%) patients reached a CAL VRS threshold of ≥0.40 whilst 54/118 (46%) reached a ≥10% FVC decline threshold (p=0.0003). 89/118 (75%) patients reached either the CAL VRS ≥0.40 or ≥10% FVC decline threshold (Table 2, Figure 4). Use of a CAL VRS threshold of ≥0.40 identified 35/118 (30%) more patients reaching an endpoint than the ≥10% FVC decline threshold alone (Table 2, Figure 4). Similarly, at least 30% more patients reached an endpoint when an UZ VRS threshold was used alongside a ≥10% FVC decline threshold (Figure 5, Supplementary Table 3).

**Figure 4a.**
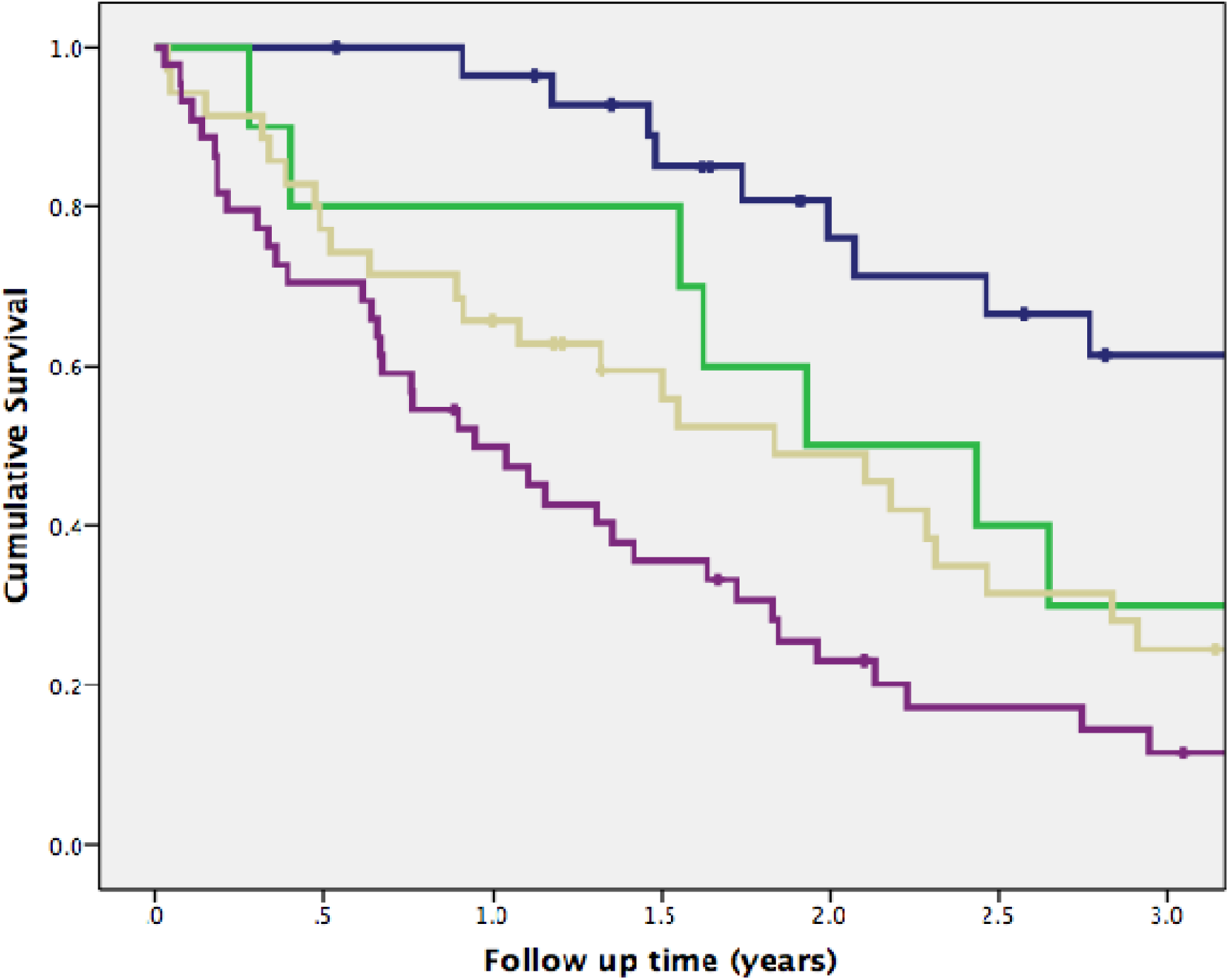
Kaplan Meier curves demonstrating 3-year survival in the IPF population not receiving antifibrotic medication. Patients were coded as undergoing an event if they had a ≥10% relative FVC decline in one year, or a VRS increase of ≥0.40 in a year. Blue=patients who did not reach either the VRS or FVC thresholds (n=29); green=patients with an FVC decline ≥10% in one year but no VRS increase ≥0.40 (n=10); yellow=patients with a VRS increase ≥0.40, but no relative FVC decline ≥10% (n=35); purple= patients with both a relative FVC decline ≥10%, and a VRS increase ≥0.40 (n=44). Log rank test=<1×10^-6^.

**Figure 4b.**
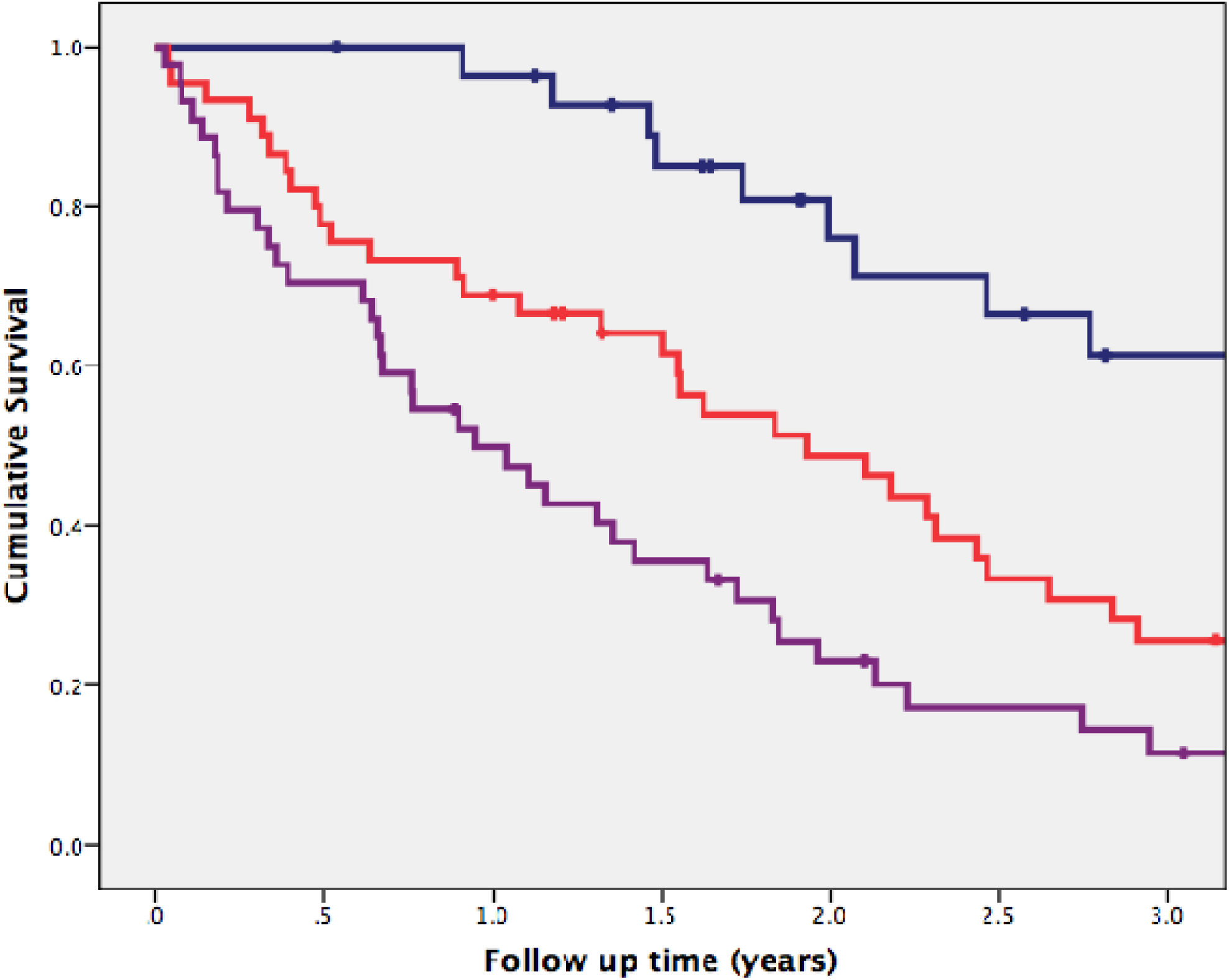
Kaplan Meier curves demonstrating 3-year survival in the IPF population not receiving antifibrotic medication. The figure is the same as Figure 4a, but patients reaching either one of the two endpoints (VRS increase ≥0.40, or FVC decline ≥10%) have been combined (red).

**Figure 5.**
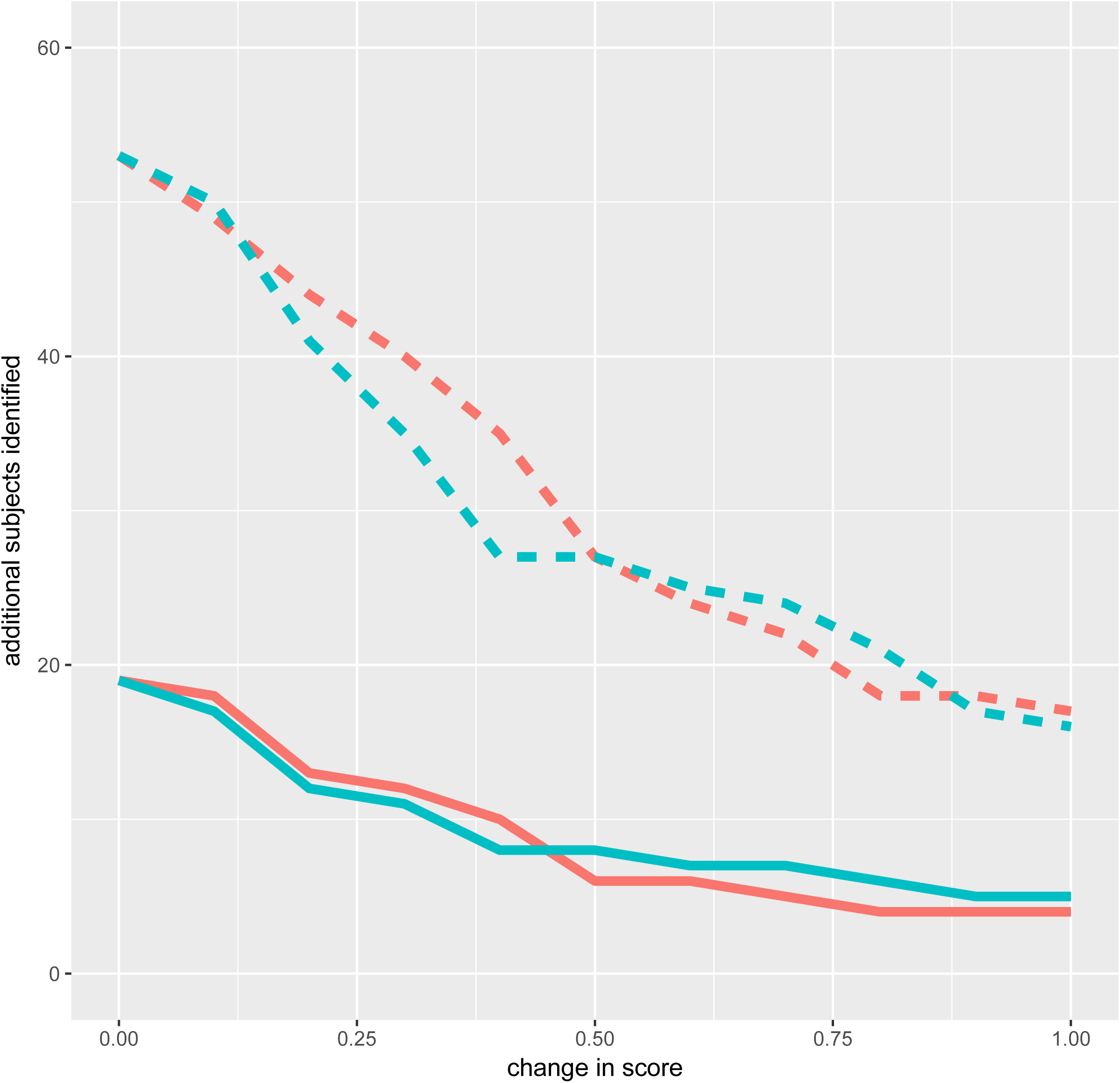
Graphs demonstrating the additional patients that would reach an endpoint (y-axis), if various vessel related structure (CAL VRS, red) or upper-zone vessel related structure (UZ VRS, green) thresholds of change (x-axis) were examined in addition to FVC decline thresholds. The FVC decline thresholds examined included a ≥5% FVC decline threshold (solid line) and a ≥10% FVC decline threshold (dotted line).

When CAL VRS and UZ VRS thresholds were examined against a ≥5% FVC decline threshold, additional patients reaching an endpoint were again identified (Figure 5, Supplementary Table 4+5). When all patients with an FVC decline more than 5% and less than 10% were subanalysed, CAL VRS thresholds ≥0.40 demonstrated C-indices that were at least equivalent to a ≥10% FVC decline threshold (Figure 6). The results suggest that a CAL VRS threshold of ≥0.40 has utility for the adjudication of marginal FVC declines between 5.0 - 9.9%, and can capture additional patients with real clinical deterioration.

**Figure 6.**
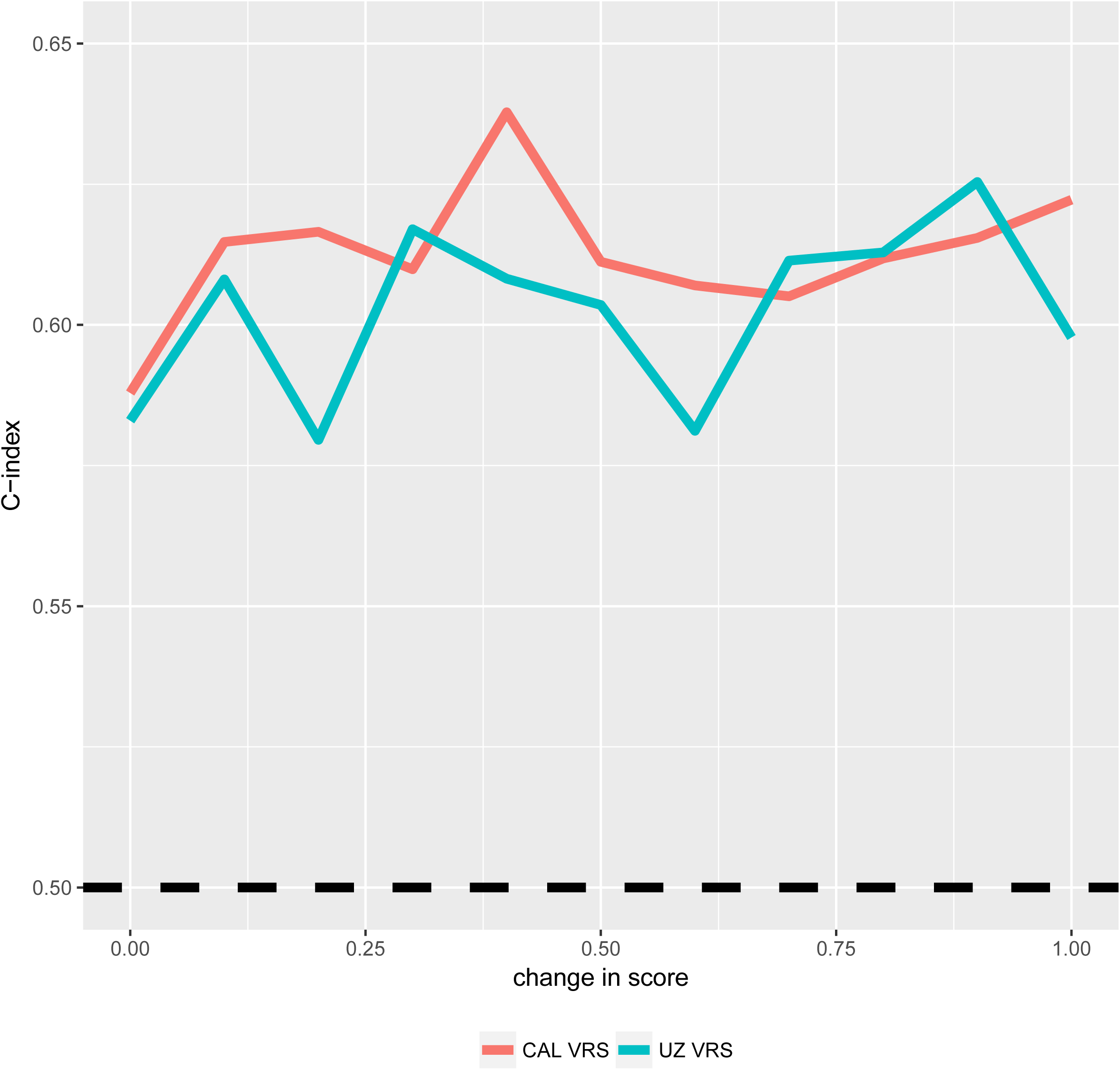
Graphs demonstrating C-indices (y-axis) for models containing varying thresholds (x-axis) of vessel related structures (CAL VRS, red) or upper-zone vessel related structures (UZ VRS, green). All models only examined patients with an FVC between 5% and 10%. The horizontal dashed black line indicates the C-index 0.5, i.e., random performance.

## DISCUSSION

Our findings demonstrate that in independent discovery and validation populations, an absolute increase in a computer-derived variable, the vessel-related structures (CAL VRS), strongly predicts mortality in IPF patients not exposed to antifibrotic medication. When FVC and CAL VRS were examined using thresholds, patients exhibiting an increase in CAL VRS ≥0.40 were not the same as those experiencing an FVC decline ≥10%. Accordingly, if a composite endpoint of CAL VRS ≥0.40 and/or ≥10% FVC decline were to be used in a drug trial setting, 30% more patients would reach the composite endpoint than with a solitary endpoint of a ≥10% FVC decline. Our findings also suggest the utility of a CAL VRS threshold ≥0.40 as an arbitration tool for marginal FVC declines (between 5.0-9.9%) to avoid the misclassification of genuine deterioration as representing measurement noise.

The link between mortality and an increase in VRS may arise from several factors. VRS may act as a surrogate marker of ILD extent. Vascular destruction in the lower lung zones in IPF, consequent to fibrosis, may steadily increase blood flow to lower pressure systems in the upper and middle zones, reflecting the mortality signal associated with an increase in UZ VRS. In addition, as fibrotic lung shrinks, spared lung containing vessels of larger size become a much greater proportion of the total lung volume, whereby VRS may act as a surrogate marker of lung volume loss. VRS may also increase as a result of traction on vascular walls secondary to the high negative intrathoracic pressure required to expand stiff fibrotic lungs. Lastly, systemic-pulmonary arteriolar shunts may develop as fibrosis progresses^12^, increasing the vascular volume in the lung.

Initiation and duration of anti-fibrotic therapy varied widely in both the discovery and validation cohorts. We therefore evaluated the prognostic significance of serial trends with the exclusion of patients taking anti-fibrotic therapy. Whilst the ideal study would examine outcomes in patients taking antifibrotic therapy throughout their disease course, it is unlikely that such a study will be possible for several years to come.

FVC has been used for over 20 years in IPF studies and has been considered the best surrogate for mortality in IPF when analysed as both a categorical and a continuous variable. Given that it is accepted that primary endpoints for IPF progression are best handled as continuous variables, and this has been the precedence in trials, we formally tested the construct validity of continuous VRS change and found that it compared favourably to FVC change in discovery and validation cohorts. When examined as a threshold, across the majority of the range of VRS change thresholds, as well as remaining independently predictive of outcome and maintaining model fit, VRS thresholds identified different poor-outcome IPF patients to FVC change thresholds. The weak correlations between FVC change and VRS change indicate that both variables represent important yet distinct surrogate measures of mortality and argues for their integration as co-endpoints rather than selecting one over another.

A ≥0·40 change in VRS across a cohort appeared to be the most accurate measure of change in VRS, when considering both its prognostic effect when judged against FVC decline and its sensitivity as an endpoint. In an individual, whilst the most accurate threshold for VRS change may also be a ≥0·40 threshold, further work is necessary to establish optimal thresholds for use in clinical practice, as just having knowledge of the range of change of a variable does not of course provide any statement of the clinical significance of that change. For example, it was noticeable that more extreme VRS cutoffs e.g. 0.75 made even more of a difference in model fit and C-index than a ≥0·40 threshold, but we cannot know how often such a magnitude of VRS change would be seen in a clinical trial population. A logical next analytic step would therefore be to evaluate VRS change in a well-controlled drug trial population receiving antifibrotics at a standardised dosing regimen.

The validity of VRS change was considered according to the OMERACT filter criteria for IPF clinical trial domains^13^. Regarding truth and discrimination criteria, VRS change was considered to be more discriminatory than FVC change at predicting outcome, with potential for use as a continuous variable (with no loss of signal strength), or as a binary threshold alongside an FVC decline threshold to improve endpoint sensitivity. The variable therefore satisfies construct and criterion validity and demonstrates sensitivity to change. The close linkages between VRS change and ILD and lung volume change satisfy face validity of VRS as a candidate endpoint. The content validity of VRS, is suggested by previous reports whereby VRS^4^ has outperformed other CT variables at predicting mortality at baseline in IPF, and zonal vessel changes have strongly predicted FVC declines in IPF^14^. Whilst computer tools are inherently reliable when examining a single scan, a degree of noise will result when CTs, reconstructed with different kernels at separate time points in the same individual are analysed. The specific impact on VRS change of differing inspiratory effort, acquisition or reconstruction parameters has not been systematically investigated, and further study is indicated. However, our analysis of this measure in a heterogeneous set of data from multiple institutions suggests this is robust. Regarding feasibility measures of VRS as an endpoint, CALIPER outputs are eminently interpretable and pose no safety issues given that they are derived from post-processing of clinically indicated scans. Accessibility and real-world utility of VRS for clinical trials or clinical practice relies on availability of repeated CTs and the computer algorithm and is therefore limited when compared to FVC measurements.

There were limitations to the current study. Though there were similar average CT intervals between the two study cohorts and change in CT variables were reported as annualized change, the CTs time intervals were not standardized in this retrospective analysis. This lack of standardization reflects real world clinical practice but may have biased our findings in patients with shorter or longer CT follow up intervals. Whilst the ideal study would have rigorous protocol-led control of serial CT and functional measurements, no such study yet exists and were it to begin today, outcome data may only be available several years hence. Accordingly, we believe our analyses capture a realistic contemporary cross-section of IPF data points. By comparing CTs at two timepoints with FVC measured multiple timepoints, we may also conceivably have punished the CT results by the introduction of noise. A naïve two-point estimate analysis was therefore performed to evaluate outcome prediction when FVC and CTs were performed at similar intervals.

In conclusion, we have demonstrated for the first time that change in a computer-derived variable, vessel-related structures, which has no visual correlate is a powerful surrogate for mortality in IPF. VRS change correlates weakly with FVC change and identifies different poor-outcome patients than a ≥10% FVC decline threshold. Use of a VRS threshold of ≥0·40 alongside a ≥10% FVC decline threshold can identify 30% more patients that reach an endpoint and argues for the consideration of VRS change as an IPF drug trial co-endpoint to adjudicate indeterminate FVC declines of 5.0-9.9%. We have also demonstrated the confounding effect on outcome of anti-fibrotic use, which will have relevance for all future outcome studies in IPF patients receiving treatment.

## Supporting information

## Authors contributions

JJ, AA, FTvB, CHMM, MV, HWvE, RC, TJ, TM, EPJ, AL, AD, TP, FM, MK, TMM, ER, AUW were involved in either the acquisition, or analysis or interpretation of data for the study. JJ and AUW were also involved in the conception and design of the study. BJB, RK and SR invented and developed CALIPER. They were involved in processing the raw CT scans and in generation of figures but were not involved with the analysis or interpretation of the data in the study.

All authors revised the work for important intellectual content and gave final approval for the version to be published. All authors agree to be accountable for the all aspects of the work in ensuring that questions related to the accuracy or integrity of any part of the work are appropriately investigated and resolved. None of the material has been published or is under consideration elsewhere, including the Internet.

## References

1. Richeldi L, du Bois RM, Raghu G, et al. Efficacy and Safety of Nintedanib in Idiopathic Pulmonary Fibrosis. N Engl J Med 2014; 370(22): 2071–82.

2. King TE, Bradford WZ, Castro-Bernardini S, et al. A Phase 3 Trial of Pirfenidone in Patients with Idiopathic Pulmonary Fibrosis. N Engl J Med 2014; 370(22): 2083–92.

3. Wells AU. Forced vital capacity as a primary end point in idiopathic pulmonary fibrosis treatment trials: making a silk purse from a sow’s ear. Thorax 2013; 68(4): 309–10.

4. Jacob J, Bartholmai B, Rajagopalan S, et al. Mortality prediction in idiopathic pulmonary fibrosis: evaluation of automated computer tomographic analysis with conventional severity measures. Eur Respir J 2016; doi:10.1183/13993003.01011-2016.

5. Jacob J, Bartholmai BJ, Rajagopalan S, et al. Predicting outcome in idiopathic pulmonary fibrosis using automated CT analysis. AJRCCM 2018: doi:10.1164/rccm.201711-2174OC.

6. Jacob J, Bartholmai B, Rajagopalan S, et al. Automated quantitative CT versus visual CT scoring in idiopathic pulmonary fibrosis: validation against pulmonary function. J Thorac Imaging 2016; 31: 304–11.

7. Bates D, Mächler M, Bolker B, Walker S. Fitting Linear Mixed-Effects Models Using lme4. Journal of Statistical Software 2015; 67: 1–48.

8. Li J, Ji L. Adjusting multiple testing in multilocus analyses using the eigenvalues of a correlation matrix. Heredity (Edinb) 2005; 95(3): 221–7.

9. Harrell FEJ. Regresssion modelling strategies. 1 ed: Springer-Verlag, New York; 2001.

10. Harrell FEJ. R package version 5.1-1.Regression Modeling Strategies: https://CRAN.R-project.org/package=rms.; 2017.

11. IBM Corp. Released 2011. IBM SPSS Statistics for Windows VA, NY: IBM Corp.

12. Turner-Warwick M. Precapillary systemic-pulmonary anastomoses. Thorax 1963; 18: 225–37.

13. Saketkoo L A, Mittoo S, Huscher D, et al. Connective tissue disease related interstitial lung diseases and idiopathic pulmonary fibrosis: provisional core sets of domains and instruments for use in clinical trials. Thorax 2014; 69(5): 436–44.

14. Jacob J, Bartholmai BJ, Rajagopalan S, et al. Serial automated quantitative C T analysis in idiopathic pulmonary fibrosis: functional correlations and comparison with changes in visual C T scores. Eur Radiol 2017; 10.1007/s00330-017-5053-z.

