## Supplementary material for "Longitudinal prediction of outcome in idiopathic pulmonary fibrosis using automated CT analysis"

**Supplementary Appendix:**

**Pulmonary Function Protocol**

Pulmonary function tests (PFT) were analysed if performed within 3 months of the corresponding CT scan according to established protocols ^1^. Spirometry (Jaeger Master screen PFT, Carefusion Ltd., Warwick, UK and Houten, NL; Puritan Bennett Renaissance pneumotachography-based flow spirometer, Mallinckrodt, St Louis, MO, USA), plethysmographic lung volumes (Jaeger Master screen Body, Carefusion Ltd., Warwick, UK and Houten, NL), and diffusion capacity for carbon monoxide (Jaeger Master screen PFT, Carefusion Ltd., Warwick. UK and Houten, NL; Puritan Bennett Renaissance pneumotachography-based flow spirometer, Mallinckrodt, St Louis, MO, USA). Parameters assessed: forced vital capacity (FVC) and single breath carbon monoxide diffusing capacity corrected for hemoglobin concentration (DLco).

**CT Protocol**

CT scans at the Royal Brompton Hospital were obtained using a 64-slice multiple detector CT scanner (Somatom Sensation 64; Siemens, Erlangen, Germany) or a 4-slice multiple-detector CT scanner (Siemens Volume Zoom; Siemens, Erlangen, Germany). All images were reconstructed using a high spatial frequency, B70 kernel (Siemens, Munich, Germany). CT scans at the St. Antonius Hospital, Nieuwegein, were obtained using a 64-slice multiple detector CT scanner (Phillips Brilliance 64; Cleveland, Ohio, USA) or a 256-slice multiple-detector iCT scanner (Phillips, Cleveland, Ohio, USA). All images were reconstructed using C, EC, L or YC kernels (Phillips, Cleveland, Ohio, USA). CT scans at the Mayo Clinic Rochester were acquired using LightSpeed Ultra (BONE kernel; GE Healthcare, Cleveland, OH, USA) and Model Sensation 64 (B46 kernel; Siemens, Munich, Germany) scanners.

All patients were scanned from lung apices to bases, at full inspiration, using a peak voltage of 120 kVp with tube current modulation (range, 30 to 140 mA). Images of 0.5-1mm thickness were viewed at window settings optimized for the assessment of the lung parenchyma (width 1500 HU; level -500 HU).

**CALIPER CT evaluation**

Data processing

Data processing steps have been previously described^2-4^. In reconstruction algorithms utilizing an edge-enhancing algorithm (Siemens B70, Phillips L and YC), a median filter was applied to 3x3x1 voxel volumes. The Siemens B80 algorithm was markedly edge-enhancing and resulted in misclassification of parenchymal features such as honeycombing. As a result, 4 patients with CT imaging reconstructed with the Siemens B80 algorithm were excluded from the study.

After the median filter pre-processing step for the other edge-enhancing CT algorithms, the lungs were extracted from the surrounding thoracic structures and then segmented into upper, middle, and lower lung zones using the carina as a landmark. Lung segmentation was performed with an adaptive density-based morphologic approach,^5^ whereas airway segmentation involved iterative 3-dimensional region growing, density thresholding (thresholds including 950HU and 960 HU) and connected components analysis. Parenchymal tissue type classification was applied to 15x15x15 voxel volume units using texture analysis, computer vision-based image understanding of volumetric histogram signature mapping features, and 3-dimensional morphology^2^.

Pulmonary vessel extraction was achieved using an optimized multiscale tubular structure enhancement filter based on the eigenvalues of the Hessian matrix. More precisely, the second-order derivatives that occurred in the regions that surrounded each pulmonary voxel were calculated and formed the Hessian matrix. The eigenvalues of this Hessian matrix were then computed and used to determine the likelihood that an underlying voxel was connected to a dense tubular structure and therefore represented a vessel or vessel-related tubular structure ^2,6^. The pulmonary VRS score (CALIPER VRS) excluded vessels at the lung hilum and expressed the VRS as a percentage of the total lung volume.

Parenchymal CT features were quantified on a total lung basis and a zonal basis. The upper and middle lung zones were demarcated at the level of the carina whilst the middle and lower lung zones were separated at the midpoint between the carina and the lowermost axial image displaying the lungs.

**Statistical Analyses**

In our primary survival analyses of patients not receiving antifibrotic medication, to compare the consistency of predictor variables, the ‑log10 p-values for each measure from the discovery and validation cohorts were plotted on the x- and y-axes, respectively. Axed were labelled using the -log10 p-value scale, and included both horizontal and vertical lines marking the Li and Ji corrected cutoff for statistical significance^8^ at 0.05^5^. There was a high degree of correlation between some tested predictors, rendering the Bonferroni corrected cutoff too strict. The Li and Ji method allows calculation of the number of effective independent tests. A Bonferroni correction was then applied to that number instead of being applied indiscriminately to all variables.

P-values based on the improved model fit of multivariate mortality models indicate the importance of a single variable, but do not convey the predictive performance of the entire model. Therefore, we calculated the C-index^9^ in the discovery and validation cohorts as well as the combined cohort of subjects not receiving antifibrotic medication. Again for completeness, we included a secondary analysis on all subjects regardless of their history of antifibrotics usage in the supplement. C-indices were based on 500 bootstrap replicates for Cox mortality models using the rms package^10^ in R.

**Characteristics of IPF patients with serial CT imaging in the discovery and validation cohorts**

| **Variable** | **Discovery Cohort** | **Validation Cohort** | **Group Comparison** |
| --- | --- | --- | --- |
| **Units: percentages unless stated** |  |  |  |
| **Median Age** | 69 | 69 | =0.90^ |
| **Male/female (ratio)** | 56/15 | 40/7 | =0.40* |
| **Mean CT interval (months)** | 1.1 ± 0.4 | 1.2 ± 0.5 | =0.25 |
| **Survival (alive/dead)** | 15/56 | 13/34 | =0.41* |
| **Baseline FVC % predicted** | 74.9 ± 22.1 | 73.6 ± 16.2 | =0.72 |
| **FVC annual decline (mls) BLUP** | 257.74 ± 135.9 | 250.1 ± 131.1 | =0.78 |
| **FVC annual decline (mls) Naive** | 298.1 ± 493.1 | 234.1 ± 382.8 | =0.42 |
| **Baseline DLco % predicted** | 39.8 ± 12.8 | 46.3 ± 12.8 | =0.009 |
| **CALIPER ILD baseline** | 21.9 ± 16.7 | 22.3 ± 15.7 | =0.89 |
| **CALIPER Fibrosis baseline** | 8.1 ± 5.6 | 7.0 ± 4.8 | =0.26 |
| **CALIPER VRS baseline** | 4.7 ± 1.7 | 5.0 ± 1.8 | =0.30 |
| **CALIPER ILD change** | 8.6 ± 17.4 | 4.5 ± 9.9 | =0.11 |
| **CALIPER Fibrosis change** | 3.5 ± 5.7 | 1.4 ± 4.1 | =0.03 |
| **CALIPER VRS change** | 1.1 ± 1.5 | 0.9 ± 1.3 | =0.42 |

Supplementary Table 1. Characteristics of IPF patients with serial CT imaging in the discovery (Royal Brompton Hospital [n=71 unless stated]) and validation cohorts (St Antonius Hospital, Nieuwegein and Mayo Clinic Rochester [n=47 unless stated]). Variables examined include: patient demographic details, pulmonary function indices and CALIPER-scored CT parameters. Data represent mean values with standard deviations unless otherwise stated. Significant differences between mean ranks of the two groups were calculated using Chi-Square test for categorical independent variables (*) and the T test for continuous variables. Significant differences between median ranks of the two groups were calculated using the Mann-Whitney U test (^). Differences in median follow up time were calculated using the Log Rank Test. FVC = forced vital capacity, DLco = diffusing capacity for carbon monoxide, ILD=interstitial lung disease, VRS=vessel-related structures.

| **FVC expression** | **Cohort** | **VRS change**  **(r value, p value)** | **UZ VRS change**  **(r value, p value)** |
| --- | --- | --- | --- |
| **FVC change**  **BLUP** | **Discovery cohort** | -0.45, 8.5x10^-5^ | -0.40, 0.0005 |
|  | **Validation cohort** | -0.31, 0.03 | -0.36, 0.01 |
|  | **Combined cohort** | -0.42, 1.8x10^-6^ | -0.40, 5.7x10^-6^ |
| **FVC change**  **Naive** | **Discovery cohort** | -0.67, 1.4x10^-10^ | -0.51, 3.5x10^-10^ |
|  | **Validation cohort** | -0.17, 0.24 | -0.22, 0.16 |
|  | **Combined cohort** | -0.53, 5.9x10^-10^ | -0.45, 1.3x10^-10^ |

Supplementary Table 2. Pearsons correlations examining linkages between forced vital capacity (FVC) change and either vessel related structure (VRS) change or upper zone vessel related structure (UZ VRS) change in IPF patients not receiving antifibrotic medication in the discovery cohort (Royal Brompton Hospital, n=71), the validation cohort (St Antonius Hospital, Nieuwegein and Mayo Clinic Rochester, n=47) and both cohorts combined (n=118). Annualised FVC change was measured using a linear mixed effects model on all eligible timepoints (best linear unbiased prediction [BLUP]), or using a naïve estimate based on two timepoints (first and last).

| **Absolute UZ**  **VRS change** | **UZ VRS=F FVC=F** | **UZ VRS=T FVC=F** | **UZ VRS=F FVC=T** | **UZ VRS=T FVC=T** |
| --- | --- | --- | --- | --- |
| **UZ VRS = 0** | 11 | 53 | 5 | 49 |
| **UZ VRS = 10** | 14 | 50 | 5 | 49 |
| **UZ VRS = 20** | 23 | 41 | 8 | 46 |
| **UZ VRS = 30** | 29 | 35 | 10 | 44 |
| **UZ VRS = 40** | 37 | 27 | 11 | 43 |
| **UZ VRS = 50** | 37 | 27 | 11 | 43 |
| **UZ VRS = 60** | 39 | 25 | 12 | 42 |
| **UZ VRS = 70** | 40 | 24 | 13 | 41 |
| **UZ VRS = 80** | 43 | 21 | 17 | 37 |
| **UZ VRS = 90** | 47 | 17 | 18 | 36 |
| **UZ VRS = 100** | 48 | 16 | 21 | 33 |

Supplementary Table 3. Numbers of patients reaching threshold endpoints of CALIPER upper-zone vessel-related structure increase (UZ VRS) or ≥10% relative forced vital capacity (FVC) decline from the study population of 118 patients not exposed to antifibrotic medication. T=true (did reach threshold), F=false (did not reach threshold).

| **Absolute CAL**  **VRS change** | **CAL VRS=F FVC=F** | **CAL VRS=T FVC=F** | **CAL VRS=F FVC=T** | **CAL VRS=T FVC=T** |
| --- | --- | --- | --- | --- |
| **CAL VRS = 0** | 4 | 19 | 13 | 82 |
| **CAL VRS = 10** | 5 | 18 | 18 | 77 |
| **CAL VRS = 20** | 10 | 13 | 19 | 76 |
| **CAL VRS = 30** | 11 | 12 | 22 | 73 |
| **CAL VRS = 40** | 13 | 10 | 26 | 69 |
| **CAL VRS = 50** | 17 | 6 | 31 | 64 |
| **CAL VRS = 60** | 17 | 6 | 35 | 60 |
| **CAL VRS = 70** | 18 | 5 | 36 | 59 |
| **CAL VRS = 80** | 19 | 4 | 40 | 55 |
| **CAL VRS = 90** | 19 | 4 | 42 | 53 |
| **CAL VRS = 100** | 19 | 4 | 45 | 50 |

Supplementary Table 4. Numbers of patients reaching threshold endpoints of CALIPER vessel-related structure increase (CAL VRS) or ≥5% relative forced vital capacity (FVC) decline from the study population of 118 patients not exposed to antifibrotic medication. T=true (did reach threshold), F=false (did not reach threshold).

| **Absolute UZ**  **VRS change** | **UZ VRS=F FVC=F** | **UZ VRS=T FVC=F** | **UZ VRS=F FVC=T** | **UZ VRS=T FVC=T** |
| --- | --- | --- | --- | --- |
| **UZ VRS = 0** | 4 | 19 | 12 | 83 |
| **UZ VRS = 10** | 6 | 17 | 13 | 82 |
| **UZ VRS = 20** | 11 | 12 | 20 | 75 |
| **UZ VRS = 30** | 12 | 11 | 27 | 68 |
| **UZ VRS = 40** | 15 | 8 | 33 | 62 |
| **UZ VRS = 50** | 15 | 8 | 33 | 62 |
| **UZ VRS = 60** | 16 | 7 | 35 | 60 |
| **UZ VRS = 70** | 16 | 7 | 37 | 58 |
| **UZ VRS = 80** | 17 | 6 | 43 | 52 |
| **UZ VRS = 90** | 18 | 5 | 47 | 48 |
| **UZ VRS = 100** | 18 | 5 | 51 | 44 |

Supplementary Table 5. Numbers of patients reaching threshold endpoints of CALIPER upper-zone vessel-related structure increase (UZ VRS) or ≥5% relative forced vital capacity (FVC) decline from the study population of 118 patients not exposed to antifibrotic medication. T=true (did reach threshold), F=false (did not reach threshold).

Supplementary Figure 1. Scatterplots demonstrating -log10 p-values for various computer-derived (CALIPER) variables (blue points) and FVC decline (yellow points) in all patients (Figure 1a) in the discovery cohort (x-axis, n=104) and validation cohort (y-axis, n=96). Horizontal and vertical dotted lines represent the Li and Ji corrected cutoff for statistical significance. FVC decline was calculated using two methods: naïve estimate from two timepoints aligned with the two CT timepoints (simple) and using best linear unbiased predictions. FVC change was expressed as a continuous variable (FVC change), and at ≥5% decline and ≥10% decline thresholds. The FVC value at the timepoint of the second CT scan (red dot) was used to benchmark expressions of FVC decline. The pulmonary vessel-related structure score (CAL VRS) was subdivided according to zonal location (UZ VRS=upper zone, MZ VRS=middle zone, LZ VRS=lower zone) and structure cross-sectional area in each zone (<5mm^2^, 5-10 mm^2^, 10-15 mm^2^, 15-20 mm^2^, >20 mm^2^).

**REFERENCES**

1.	Quanjer PH. Standardized lung function testing. *Eur Respir J - Suppl* 1993; **6**: 1-100.

2.	Bartholmai BJ, Raghunath S, Karwoski RA, et al. Quantitative CT imaging of interstitial lung diseases. *J Thorac Imaging* 2013; **28**(5): 298-307.

3.	Maldonado F, Moua T, Rajagopalan S, et al. Automated quantification of radiological patterns predicts survival in idiopathic pulmonary fibrosis. *Eur Respir J* 2014; **43**(1): 204-12.

4.	Jacob J, Bartholmai B, Rajagopalan S, et al. Automated quantitative CT versus visual CT scoring in idiopathic pulmonary fibrosis: validation against pulmonary function. *J Thorac Imaging* 2016; **31**: 304-11.

5.	Hu S, Hoffman EA, Reinhardt JM. Automatic lung segmentation for accurate quantitation of volumetric X-ray CT images. *IEEE Trans Med Imaging* 2001; **20**(6): 490-8.

6.	Shikata H, McLennan G, Hoffman EA, Sonka M. Segmentation of pulmonary vascular trees from thoracic 3D CT images. *Int J Biomed Imaging* 2009: 11.
