## Supplementary figures and images for "Longitudinal prediction of outcome in idiopathic pulmonary fibrosis using automated CT analysis"

### Supplementary file 2

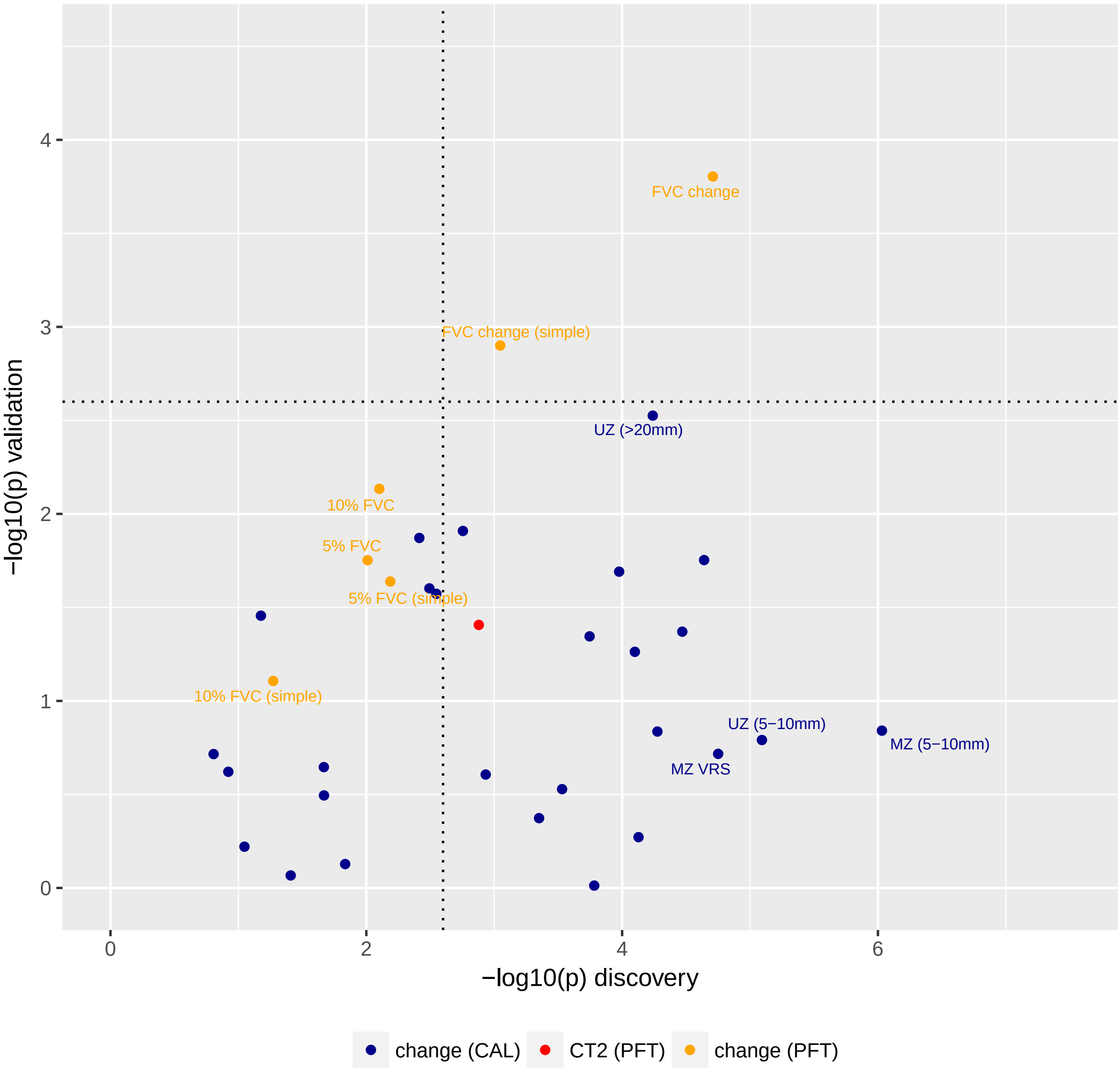
